# Prediction of therapy outcomes of CLL using gene expression intensity, clustering, and ANN classification of single cell transcriptomes

**DOI:** 10.1101/2021.08.08.455551

**Authors:** Minjie Lyu, Huan Jin, Anthony Bellotti, Xin Lin, Zhiwei Cao, Derin B. Keskin, Vladimir Brusic

**Affiliations:** School of Computer Science, University of Nottingham Ningbo China; School of Life Sciences and Technology, Tongji University, Shanghai, China; Translational Immuno-Genomics Lab, Dana-Farber Cancer Institute, Harvard Medical School, Boston, USA

**Keywords:** artificial neural networks, chronic lymphoid leukemia, gene expression counts, Ibrutinib, single cell transcriptomics, supervised machine learning

## Abstract

**Background:** Single cell transcriptomics is a new technology that enables us to measure the expression levels of genes from an individual cell. The expression information reflects the activity of that individual cell which could be used to indicate the cell types. Chronic lymphocytic leukemia (CLL) is a malignancy of B cells, one of the peripheral blood mononuclear cells subtypes. We applied five analytical tools for the study of single cell gene expression in CLL course of therapy. These tools included the analysis of gene expression distributions – median, interquartile ranges, and percentage above quality control (QC) threshold; hierarchical clustering applied to all cells within individual single cell data sets; and artificial neural network (ANN) for classification of healthy peripheral blood mononuclear cell (PBMC) subtypes. These tools were applied to the analysis of CLL data representing states before and during the therapy.

**Results:** We identified patterns in gene expression that distinguished two patients that had complete remission (complete response), a patient that had a relapse, and a patient that had partial remission within three years of Ibrutinib therapy. Patients with complete remission showed a rapid decline of median gene expression counts, and the total number of gene counts below the QC threshold for healthy cells (670 counts) in 80% of more of the cells. These patients also showed the emergence of healthy-like PBMC cluster maps within 120 days of therapy and distinct changes in predicted proportions of PBMC cell types.

**Conclusions:** The combination of basic statistical analysis, hierarchical clustering, and supervised machine learning identified patterns from gene expression that distinguish four CLL patients treated with Ibrutinib that experienced complete remission, partial remission, or relapse. These preliminary results suggest that new bioinformatics tools for single cell transcriptomics, including ANN comparison to healthy PBMC, offer promise in prognostics of CLL.

## Background

Cancer is a group of highly heterogeneous diseases characterized by abnormalities of cells (deregulation, mutation, and genome instability) that sustain immortality, uncontrolled growth, spread, and evasion of the defense mechanisms in the host organism [1]. Gene expression analysis (transcriptome analysis) was used to classify, discover subtypes, and predict outcomes in cancer [2,3]. Most gene expressions studies use analysis of bulk samples that measure average values of gene expression across all cells in the sample. Such studies can return useful information about gene expression differences between samples, but they often miss variability at the individual cell level [4]. Even within one cancer, cancer cells show very high heterogeneity reflected in differences in cellular morphology and phenotypes, including gene expression profiles, metabolic properties, motility, growth, and spread capabilities [5]. Heterogeneity of cancer cells reflects the diversity of phenotypes that differentially promote cancer progression, metastasis and resistance to therapies making clinical applications difficult [6]. The diversity of cellular and molecular phenotypes within cancer remains a crucial challenge for clinical applications including diagnosis, classification, prognosis, and therapy selection. Heterogeneity within a cancer is driven by genetic, systemic, and environmental triggers whose interplay shapes phenotypes related to cancer [7].

Single cell transcriptomics (SCT), also known as scRNA-seq, is an emerging technology that enables gene expression profiling at the individual cell level. SCT data are high-dimensional – more than 30,000 features are measured from single cells. A single representative data file may have 108 data points in form of sparse matrices where 95-99% of gene expression counts are zeros [8,9]. SCT produces Big Data – more than 20,000 data sets have been generated since 2017 and are available to public. The availability of these huge and rapidly increasing data sets requires significant data analysis efforts. The SCT data analysis or “bioinformatics pipelines” has upstream and downstream components. The upstream component is concerned with accurate and reproducible quantification of gene expression. It involves two principal steps: quality control (reads, mapping, and cell quality control), and quantification (transcripts quantification, normalization, and control of confounding factors) [10]. SCT methods show high reproducibility of gene expression profiles from samples that are prepared using standard operating procedures (manufacturer’s protocol) and identical sample processing conditions [11]. However, even relatively small changes such as updating reagent kits, for example chemistry v2 *vs*. v3 for 10x Chromium (support.10xgenomics.com) produce marked changes in gene expression profiles. We found that gene expression profiles of healthy peripheral blood mononuclear cells (PBMC) of two samples from different individuals generated from the same chemistry are more similar than the profiles from the samples from one individual generated by different chemistry [12].

The design of SCT experiments has key variables that affect the results of upstream data analysis and change gene expression profiles. These variables include the number of cells that are sequenced, cell isolation methods, experimental protocol, inclusion of quantitative standards, sequencing depth, and technical factors (instrument settings, chemistry, library versions, and software versions) [10]. The downstream analysis of SCT data is still developing, and significant challenges are still unresolved. The challenges for the downstream SCT data analysis include [13]: handling sparsity in SCT data; lack of data-analytical tools for analysis of differential gene expression; lack of practically useful reference data sets; dealing with dynamic changes in cells over time, state, and space; handling errors and missing data; dealing with heterogeneity in tumors; integration of SCT data across samples, experimental designs, and protocols; and validation and benchmarking of SCT data analysis tools. Because of the sparseness of SCT data matrices and the lack of representative reference data sets, classification of single cells and differential gene expression analysis typically deploys unsupervised methods to assist cell type assignments (cell classification) before cell differential analysis of gene expression can be performed. Unsupervised clustering is currently the method of choice for classification of cell types and subtypes [14–15]. Most SCT data analyses start with unsupervised clustering followed by annotation of clusters and cells using various tools [16]. Cell labelling usually involves the analysis of canonical markers. This process is inefficient and involves manual steps – the resulting models display unstable accuracy and poor generalization between different studies [13,15,17,18]. Supervised machine learning methods address the discussed limitations by learning the properties of cell classes from labelled reference data sets using model training. The model is then applied to unlabelled data to annotate each individual cell. This learning is done through the inclusion of representative data sets and model validation [9,19]. A selection of supervised methods has been proposed for the classification of cell types and subtypes [18–23]. These methods are promising but due to the lack of comprehensive representative data sets these methods are yet to be validated using multiple independent data sets.

The challenge for the analysis of SCT data in cancer is our ability to deal with the heterogeneity of cells, both between cancers (population studies), within a cancer (individual disease), and deal with data sparseness and technical factors. The key SCT tasks include identification of cancer and non-cancer cells, understand cancer microenvironment, assess the effects of therapies, understand dynamic changes in cancer cells, cancer microenvironment, and the organism [24]. For example, single cell transcriptomics offers insight into the cancer microenvironment [25], drivers of metastasis [26], and the effects of cancer therapy [27]. Furthermore, the analysis of 123 PBMC cell subtypes in individuals shaw changes in numbers, mostly decreases, with age (12 cell subtypes show significant changes, p<0.05), and increases with metastatic cancer (23 cell types, p<0.05) [28]. SCT has excellent potential to revolutionize diagnosis, classification, therapy selection, and prognosis in cancer, but there is an urgent need for tools and methodology that will ensure reproducibility of SCT results. These tools include comprehensive reference data sets, standardized sample processing procedures, and supervised machine learning methods for cell classification.

We defined a diverse reference data set for classification of healthy PBMC and trained an artificial neural network (ANN) to achieve 95% accuracy in 5-class classification of PBMC [29]. The five main cell classes in PBMC include B cells (BC), dendritic cells (DC), monocytes (MC), natural killer cells (NK), and T cells (TC). In another study, we trained an ANN to assess the progress of chronic lymphocytic leukemia (CLL) upon therapy by kinase inhibitor Ibrutinib. The task was to classify cells into pre- and post-treatment groups using SCT gene expression changes by [30]. The study results indicated that the changes in gene expression at 30 days after the start of therapy could predict cancer responses to the therapy.

We used the same standardized format of the SCT matrices and trained artificial neural networks for classifying health PBMC subtypes to classify PBMC in CLL samples before and during the Ibrutinib treatment. We pursued three research questions in this study:

- How do SCT profiles in CLL patients change during the ibrutinib therapy?
- What are the differences in single cell gene expressions between healthy and CLL subjects?
- Can we accurately predict the outcome of Ibrutinib therapy and how early these predictions can be made?

## Methods

This study was performed using data from previously published CLL study [27] of four patients that were treated by Ibrutinib, an irreversible Bruton’s tyrosine kinase (BTK) inhibitor. Ibrutinib binds to BTK with high affinity and inhibits B-cell receptor signalling. This inhibition results in decreased B-cell activation, proliferation, and cell survival. Ibrutinib demonstrates anti-tumor activity in B-cell cancers [31]. Our study focuses on the analysis of SCT data from four patients treated by Ibrutinib (two with full remission, one with partial remission, and one with relapse). For this analysis we used SCT classification of healthy PBMC by ANN that we developed earlier [29] to assess the SCT changes associated with successful CLL treatment.

### Study design

In our previous study, we determined that patterns of gene expression in patients with CLL differ between samples taken before the treatment and at time points during the therapy. We trained an ANN with data sets from different time points and predicted PBMC cells on “Day 0” (before therapy) and “Day 30”, “Day 120”, “Day 150”, and “Day 280” (during therapy)” [30]. We demonstrated that ANN distinguishes PBMC before and during treatment in successful therapy. The current study is the extension of our studies where we trained ANN to classify PBMC in five main classes. Our trained network reported 95% accuracy in classification of healthy PBMC into 5 main classes (positive predictive values were BC – 92.0%, DC – 94.3%, MC – 97.0%, NK – 81.5%, and TC – 97.4%) [29]. We applied 5-class classification to CLL PBMC samples to observe the behavior of PBMC in CLL before and during therapy. The overall study design is shown in Figure 1. It has four main steps: data pre-processing and standardization, basic statistical analysis of data sets (statistical distributions of gene expression counts), followed by the analysis of changes of counts during treatment. We then performed hierarchical clustering of all cells in each CLL file to check their similarities and groupings. Lastly, we performed the prediction of cell types by our trained ANN model and interpreted the results.

**Figure 1.**
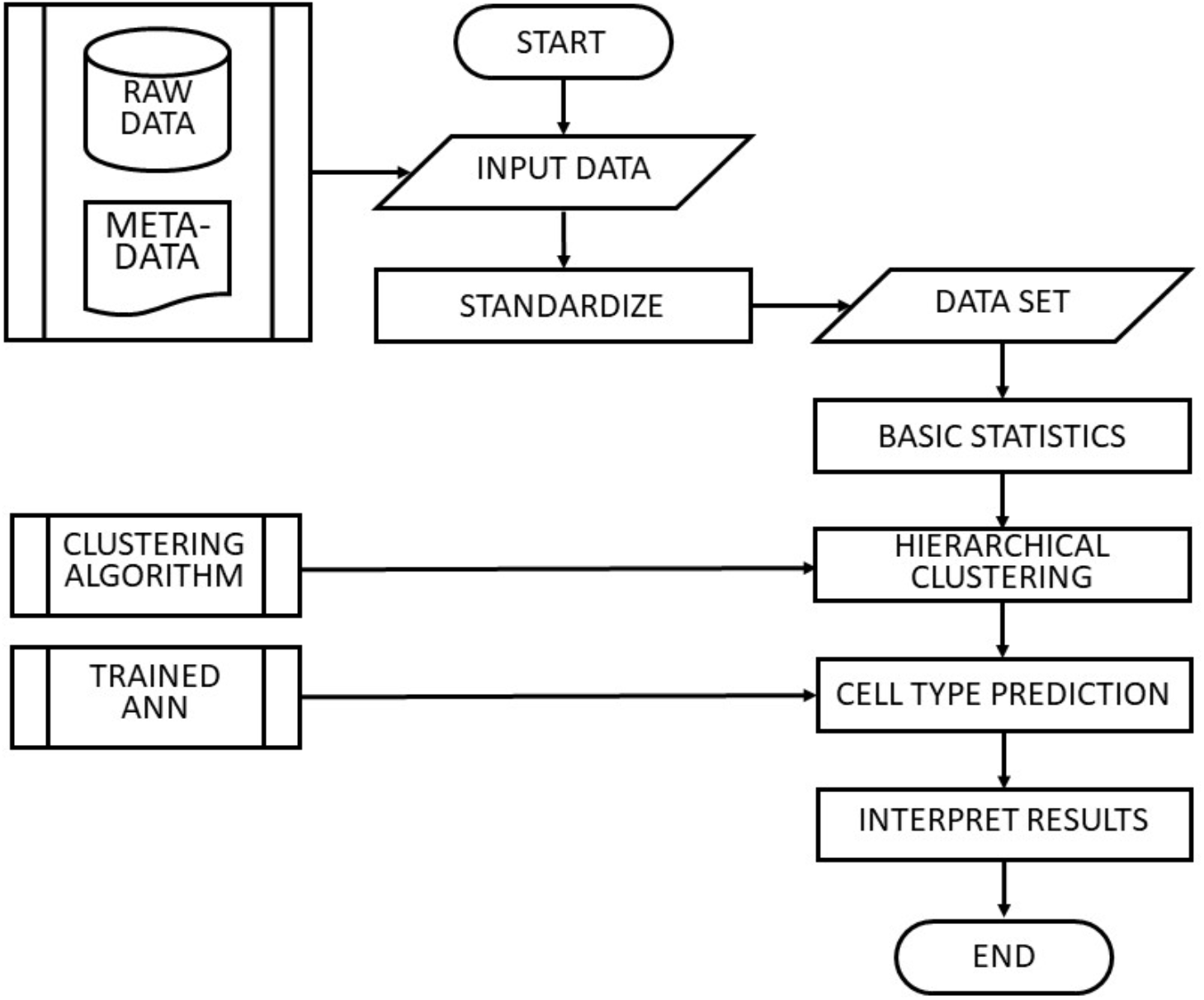
The design of this study. Raw data and metadata were obtained from [27]. Standardization of data sets was performed as described earlier [9]. The ANN model was developed as described in [9,29]. The details of the study are further elaborated in the main text.

### Data sets

Data were extracted from the NCBI Gene Expression Omnibus (GEO) database (www.ncbi.nlm.nih.gov/geo). The GEO accession of the study is GSE111014. The IDs of the 12 samples are shown in Table 1. The description of the study has been reported in [27]. PBMC was obtained from 4 CLL patients (CLL1, CLL5, CLL6, and CLL8) before treatment (day 0) and on treatment days 30 and 120. CLL5 had a sample collected on day 150 instead of day 120, and CLL6 had an additional sample taken on day 280. The number of datasets indicating cell numbers in each sample is shown in Table 2. The total number of cells we used in this study is 48,016. The breakdown of cell numbers by sample collection days is shown in Table 3. CLL1 and CLL8 experienced full remission, CLL6 had partial remission, while CLL5 had the relapse of CLL.

**Table 1.**
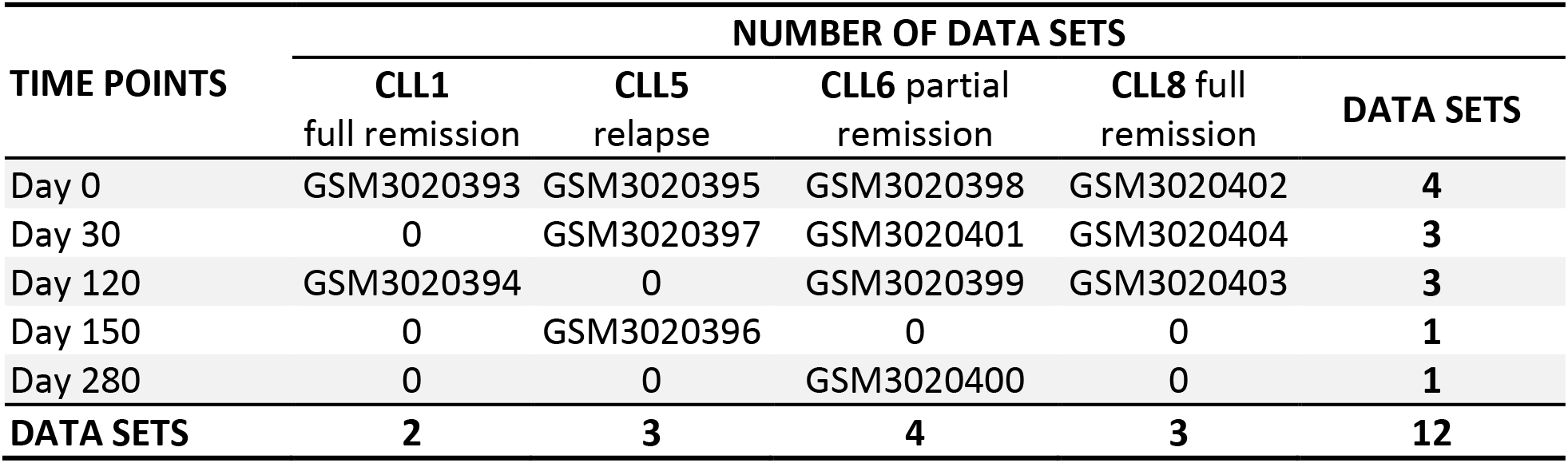
Twelve data sets used in this study were obtained from the GEO database (www.ncbi.nlm.nih.gov/geo) from study GSE111014 [27]. They represent SCT data at several therapy time points (Day 0 is before therapy) for four patients (CLL1, CLL5, CLL6, and CLL8).

**Table 2.**
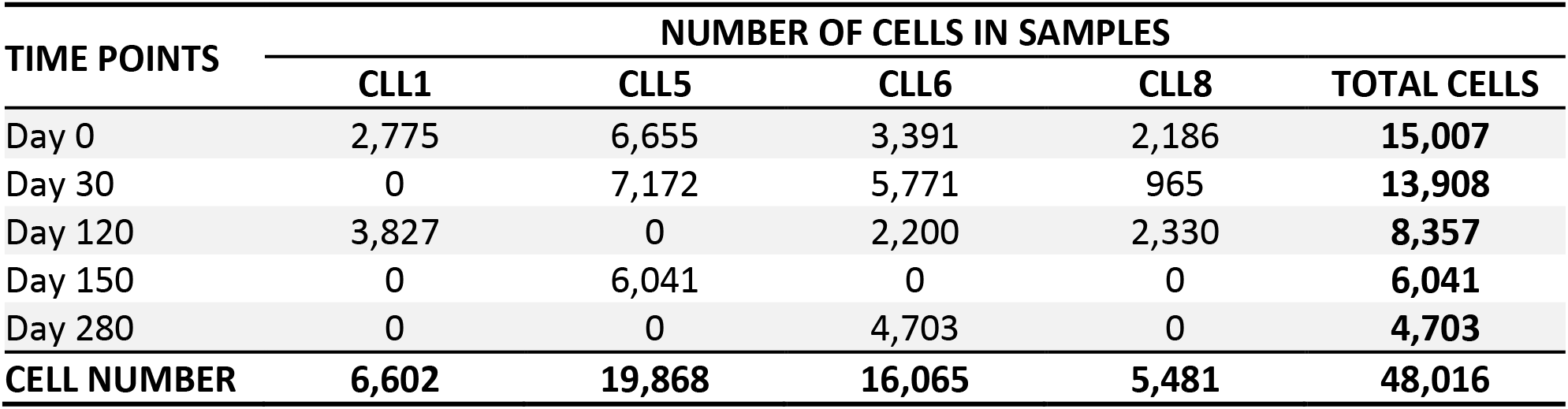
The number of cells from samples by patients and therapy time points.

**Table 3.** The values of median gene counts and interquartile (IQR) ranges of data sets representing samples taken from individual patients on specific days. FR – full remission, PR – partial remission, RE – relapse.

| PATIENT | Day 0 |  | Day 30 |  | Day 120 /150* |  | Day 280 |  |
| --- | --- | --- | --- | --- | --- | --- | --- | --- |
|  | MEDIAN | IQR | MEDIAN | IQR | MEDIAN | IQR | MED | IQR |
| CLL1 (FR) | 1922 | 1241 | na | na | 161 | 271 | na | na |
| CLL8 (FR) | 2315 | 1222 | 527 | 548 | 307 | 338 | na | na |
| CLL6 (PR) | 2060 | 1045 | 1762 | 721 | 1615 | 785 | 529 | 299 |
| CLL5 (RE) | 1483 | 822 | 1920 | 640 | 1095* | 409* | na | na |
| Healthy control | 1397 | 340 |  |  |  |  |  |  |

The metadata describing the experiment is shown in Supplemental Table 1. Each file was analysed for the distribution of total feature (gene expression) counts. The distributions of gene expression values across cells in each data set were visualized. We used seaborn [32] module seaborn.violinplot to draw the violin plots of feature distributions, and “area” as the value for the parameter “scale” to ensure that all violin plots in our analysis have the same area. The violin plots with the same area show the changes and different in gene expression level between multiple therapy points and patients. For the analysis of healthy PBMC data sets, we used quality control (QC) thresholds to eliminate cells that have less than 670 total feature counts or less than 300 positive features. We deployed this threshold on CLL data with intention to eliminate cells of presumably low quality. However, we found that the in samples from treatment period for CLL1 and CLL8 (full remission), as well as the last sample from CLL6 (partial remission) had both very low total counts and low positive feature numbers, below the QC thresholds (Supplemental Table 2). And the high percentage of cells above the QC thresholds before the therapy for all 4 patients gave the confidence that low percentage above the QC threshold after therapy was not caused by experiment fact. We then performed comparative analysis of the total counts to identify possible correlations with treatment outcomes using comparison of the percentage of cells that were above the QC threshold. The percentage of cells above the QC corresponds to the overall activity of cells in the studied sample.

### Classification of cells from CLL samples

The main characteristic of CLL is an extensive clonal expansion of cancerous B cells, and profound changes in the regulation of immune cells [33]. In CLL, the number of B cells increase in numbers by approximately 100-fold (as compared to healthy status), T cells double their number, NK cells remain similar, while the number of monocytes drops by 5-fold. Ibrutinib therapy reverses the composition of PBMC cell types towards healthy reference ranges [33]. To understand the similarity of cells within and between cell types in CLL samples, we performed cell similarity analysis using hierarchical clustering. We used the function ‘clustermap’ from the Seaborn [32] Python library, as described in [34]. A sparse matrix is taken as input data for the seaborn.clustermap() function to calculate a heatmap of hierarchically clustered elements (gene expression profiles of each cell). The input parameters were assigned: linkage – average (UPGMA algorithm), metric (distance metric) – correlation representing Pearson correlation distance range from −1 to 1). Other parameters in the clustermap() function assumed their default values. Heatmaps were produced for all twelve CLL files described in Table 1 and for the healthy control data set BroadS2 (Supplemental Table 1). The heatmaps were compared for each patient along the timeline (between patients, and between CLL therapy time and healthy control data set). The heatmaps of hierarchical clustering show how cells group together and the similarity of them (range 0 to 1, where 1 represents identity), and also the different and change between patients and multiple therapy points.

Next, we performed the classification of cells from each CLL sample to assess the changes of proportions of each cell type before and during the ibrutinib therapy. We used ANN reported in [29] that has 30,698 input units, ten hidden layer units, and five output units. The activation function was rectified linear unit and solver was Adam (adaptive moment estimation) (Figure 2). This trained ANN was used to assign the 5 main PBMC subtype to the CLL samples before and during the therapy. The ANN was assessed as 95% accurate in classification of five PBMC cell types (BC, DC, MC, NK, and TC) [29]. We used radar plots to compare different patients on the same therapy day, and also compared changes of cell type proportions for each patient, over time. DC were excluded from further analysis because their absolute counts of DC in CLL are unknown and, because of their low frequency (around 1.5% of the total PBMC) in CLL samples DC frequencies are very close to zero, so these results were not informative. Although the prediction of cell types in CLL patients are not necessarily true cell types, the change and different proportion of each cell type between therapy points and patients reflect the effect of the Ibrutinib.

**Figure 2.**
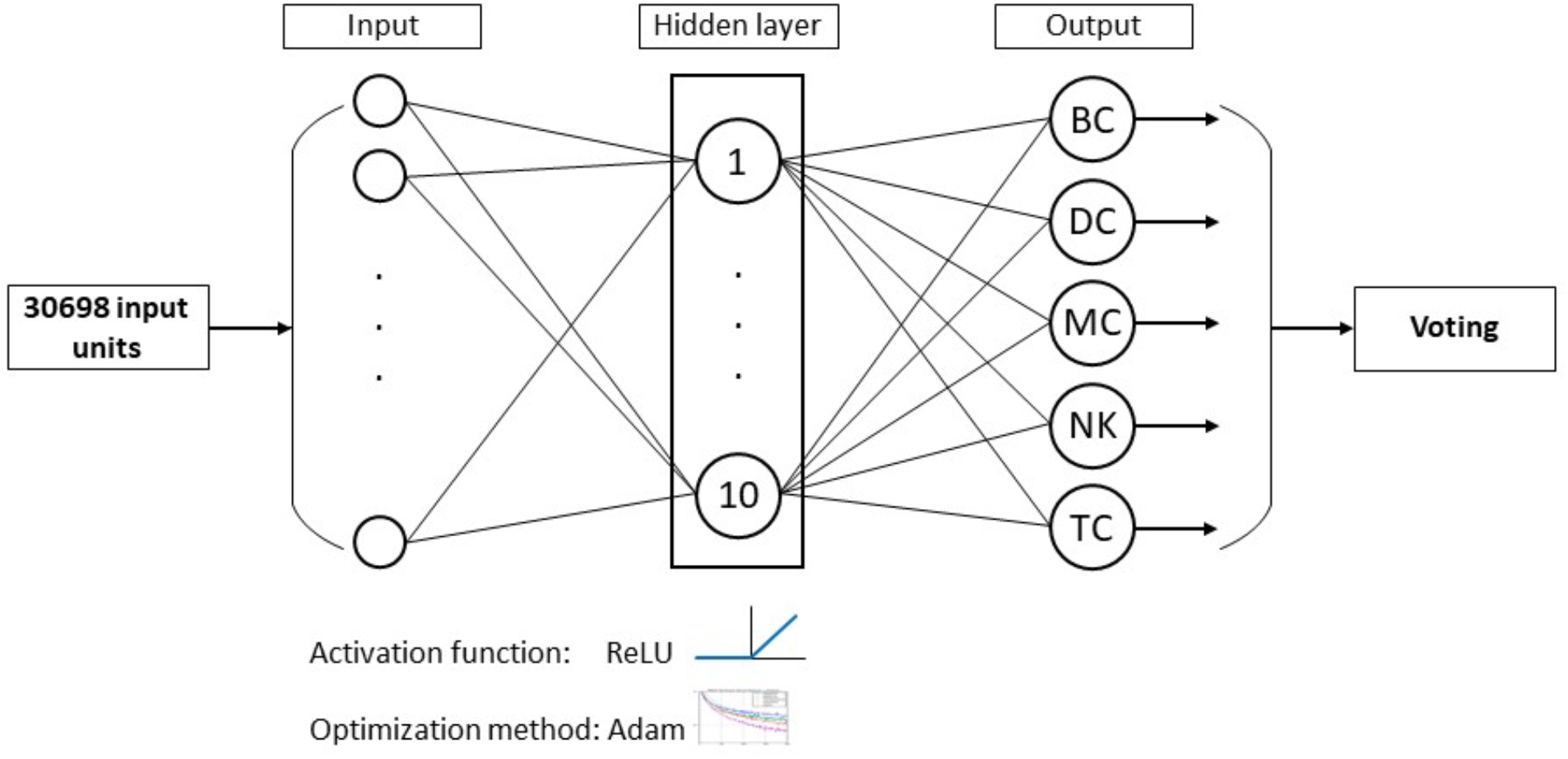
Artificial Neural Network architecture. The 30,698 input units represent our standardized list of features (genes). The output nodes provide signal values that predict cell class: B cells (BC), dendritic cells (DC), monocytes (MC), natural killer cells (NK), and T cells (TC). The final label of each query cell is determined by 10 cycles of voting.

## Results

### Statistical analysis of the CLL data sets

In comparison to data set from healthy PBMC violin plots that has leaf-like shape, moderate and narrow interquartile range (IQR), pre-therapy CLL PBMC have elongated, needle-like shape with high median value of counts and broad IQR (Figure 3 and Table 3). Patients CLL1 and CLL8 whose outcome was complete remission, showed rapid changes in their total count distributions: median distribution and the range of values of total counts rapidly declined to a very low level (their median values were lower than our standard QC threshold). Patient CLL6 whose outcome was partial remission showed no changes in total count median and the range of values during the first 120 days of Ibrutinib therapy. Only at treatment day 280, the violin plot showed low median value of total counts and a narrow IQR. Patient CLL5 whose outcome was relapse, showed an increase of total counts median and reduction of IQR at day 30 and the reduction of both measures at therapy day 150 but these values were above our standard QC threshold (670 counts). The median value of counts in day 0 of CLL1, CLL6, and CLL8 were higher than the healthy control by 40-65%, while CLL5 had similar number of total counts as healthy control. On the other hand, all CLL samples had larger IQR ranges than healthy control (140-265% increase relative to the healthy control).

**Figure 3.**
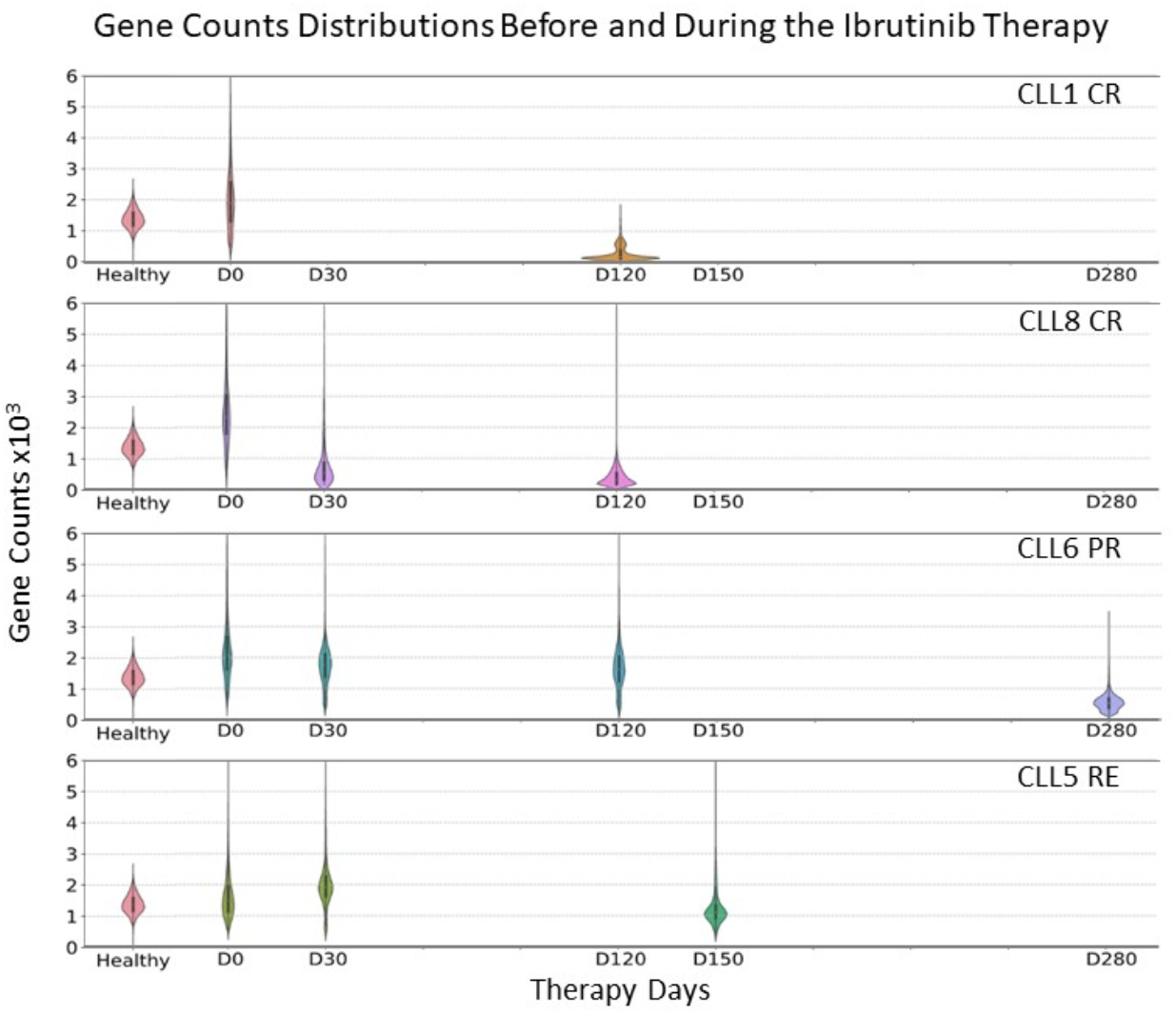
Violin plots show statistical distributions of the total counts in each data set. Healthy control (BroadS2) dataset is used as a reference. All plots have the same area size (seaborn.violinplot parameter “scale” value was set to “area”), allowing a direct comparison of total counts distribution. Patients CLL1 and CLL8 (complete remission outcome) show rapid reduction of the number of total counts with very low median after 120 days. Patient CLL6 (partial remission outcome) showed little change in total counts distribution during the first 120 days but showed a marked reduction of median gene counts at treatment day 280. Patient CLL5 (relapse outcome) showed different patterns of total counts to the patterns seen in the other three patients.

We observed the pattern where patients who experienced complete remission (CLL1 and CLL8) had a fast reduction of both median values and IQR of total counts (Fig. 4 and Supplemental Table 2). The decline of the number of total gene expression counts across all cells were observed in samples from patients that had remission as treatment outcome. The rapid decline of total counts was consistent with full CLL remission, while it was delayed in partial remission. Rapid decline of total counts was not observed in patient CLL5 that had CLL relapse as the outcome.

**Figure 4.**
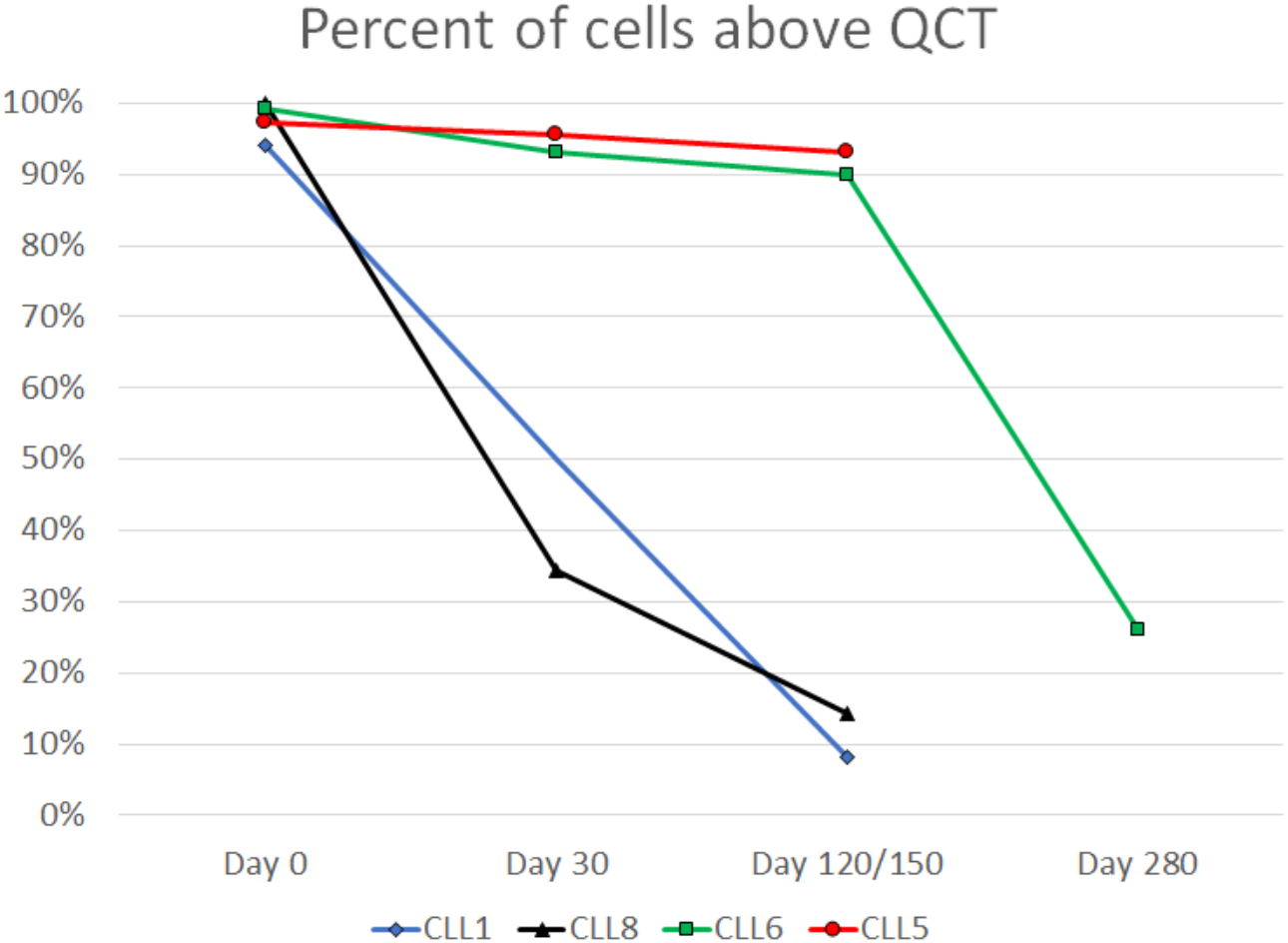
Plot of the percentage of cells ≥ QCT (greater or equal than standard QC threshold of 670 total counts) *vs.* therapy day. CLL1 and CLL8 had full remission, CLL6 had partial remission, and CLL8 had relapse, as outcomes. Patients CLL1 and CLL8 showed a rapid decline of total gene expression counts (15% or less cells had total gene expression counts ≥ 670) by the therapy day 120. In the same period, the other two patients showed minor reduction of the total gene expression counts (90% or more cells had total gene expression counts ≥ 670). Patient CLL6 showed decline in total gene expression counts between days 120 and 280, consistent with partial remission. Patient CLL5 who had relapse, showed minimal reduction in the total number of gene expression counts up to therapy day 150.

### Hierarchical clustering

Hierarchical clustering was performed on 13 files using Pearson coefficient correlation to discover groupings in the SCT data of each sample. These groupings are formed by the algorithm that analyses individual single cell gene expression profiles (containing 30,698 features). The cells that have broadly similar gene expressions profiles are grouped together. CLL is characterised by clonal expansion of malignant B cells [33] and is subject to a profound dysregulation of immune cells from both innate and adaptive branches of immune function resulting in profound changes in populations of dendritic cells monocytes, NK cells, and T cells [35].

Hierarchical clustering of twelve CLL data sets and healthy control showed major differences in cluster maps (heatmaps), both between healthy and CL L and between pre-treatment and during-treatment (Figure 5). The cluster map of a healthy sample (BroadS2) showed a large diversity of groups with tens of mini clusters showing mostly moderate mutual similarity, between 0.4 and 0.6 (Figure 5). CLL cluster maps before treatment show a smaller number of clusters than healthy samples that show high inter-cluster similarity (mostly 0.5-0.9), with marginal changes on day 30 of the treatment. Cluster maps on day 120 (CLL1, CLL8), and day 150 (CLL5) show visible changes in diversification of clusters. This diversification and reduction of inter-cluster similarity is pronounced in CLL1 day 120 but is also visible in CLL8. At therapy day 120, CLL5 shows the reduction of similarity within the main cluster region emergence of another cluster different from primary CLL (lower right corner of the cluster map), but with little diversity within the new emerging cluster. The CLL6 sample cluster maps show changes of cluster map similarities, even the overall increase of inter-cluster similarity on day 30 and 120, as compared to day 0. Marginal reduction of inter-cluster is visible on treatment day 280. The most exciting finding is that CLL1 started developing a healthy-like structure by day 120 (the bottom-right section of the cluster map).

**Figure 5.**
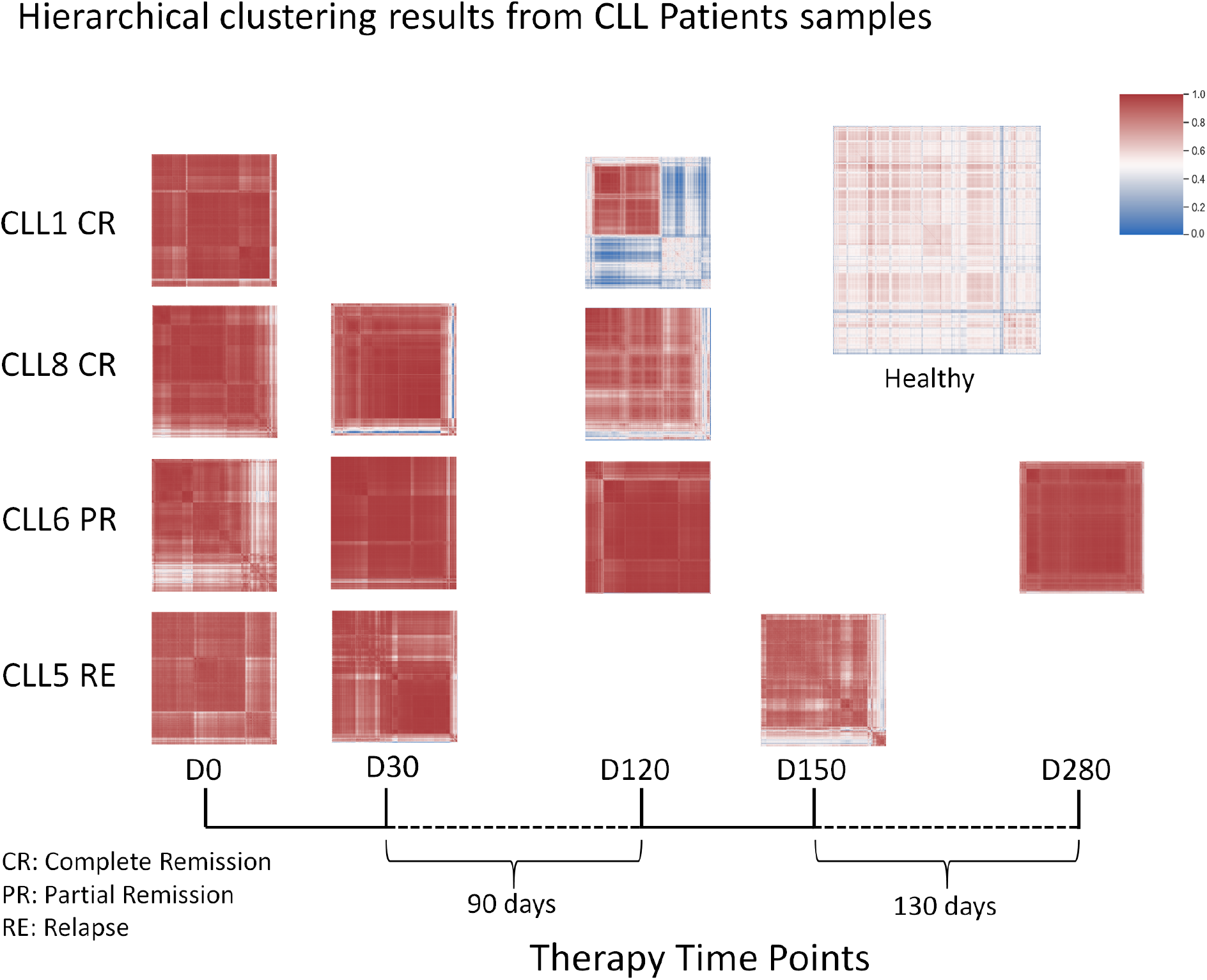
Cluster maps of CLL and a healthy control data set. Cluster maps are aligned by time (treatment days D0-D280) and patient (CLL1, CLL8, CLL 6, and CLL5, arranged by clinical outcomes).

### Prediction of similarity to healthy PBMC cell types by an ANN model

We used the predictor of healthy PBMC to assess the similarity to healthy PBMC of CLL patients during the course of therapy. In response to Ibrutinib, CLL1 had complete remission and moderate lymphocytosis (an increase of the number of lymphocytes), CLL8 had complete remission and weak lymphocytosis, CLL5 had relapse after BTK mutation and fast lymphocytosis, while CLL6 had partial remission and extended lymphocytosis.

We used ANN predictor that classifies healthy PBMC with high accuracy (95% in 5-class classification). The classifier tells us how similar gene expression of each cell is to the profiles of healthy PBMC cell types and quantifies the similarity of each cell to the healthy PBMC profile. The rough estimates of cell type content in healthy human PBMC are 5-15% BC, 1-2% DC, 10-30% MC, 5-10% NK, and 40-70% TC [9,36]. Our control data set BroadS2 is composed of 15.33% BC, 2.2% of DC, 17.34% MC, 6.85% NK, and 58.28% TC.

#### *Therapy Day 0* (Figure 6)

Predicted B cells percentages were elevated in CLL5 (81% of the total), CLL1 (67%), and CLL8 (45%) while in CLL6 (16%) they were comparable to healthy control (15%). Predicted percentages of T cells were low in CLL5 (18% of the total) and CLL1 (33%), low-normal in CLL8 (48%), and high-normal in CLL6 (70%). Predicted percentages of monocytes were low in CLL1 (0%), CLL5 (0%), CLL8 (5%), and normal in CLL6 (13%). Predicted percentages of NK cells were low in all patients – 2.5% in CLL8 and below 0.5% in other patients.

**Figure 6.**
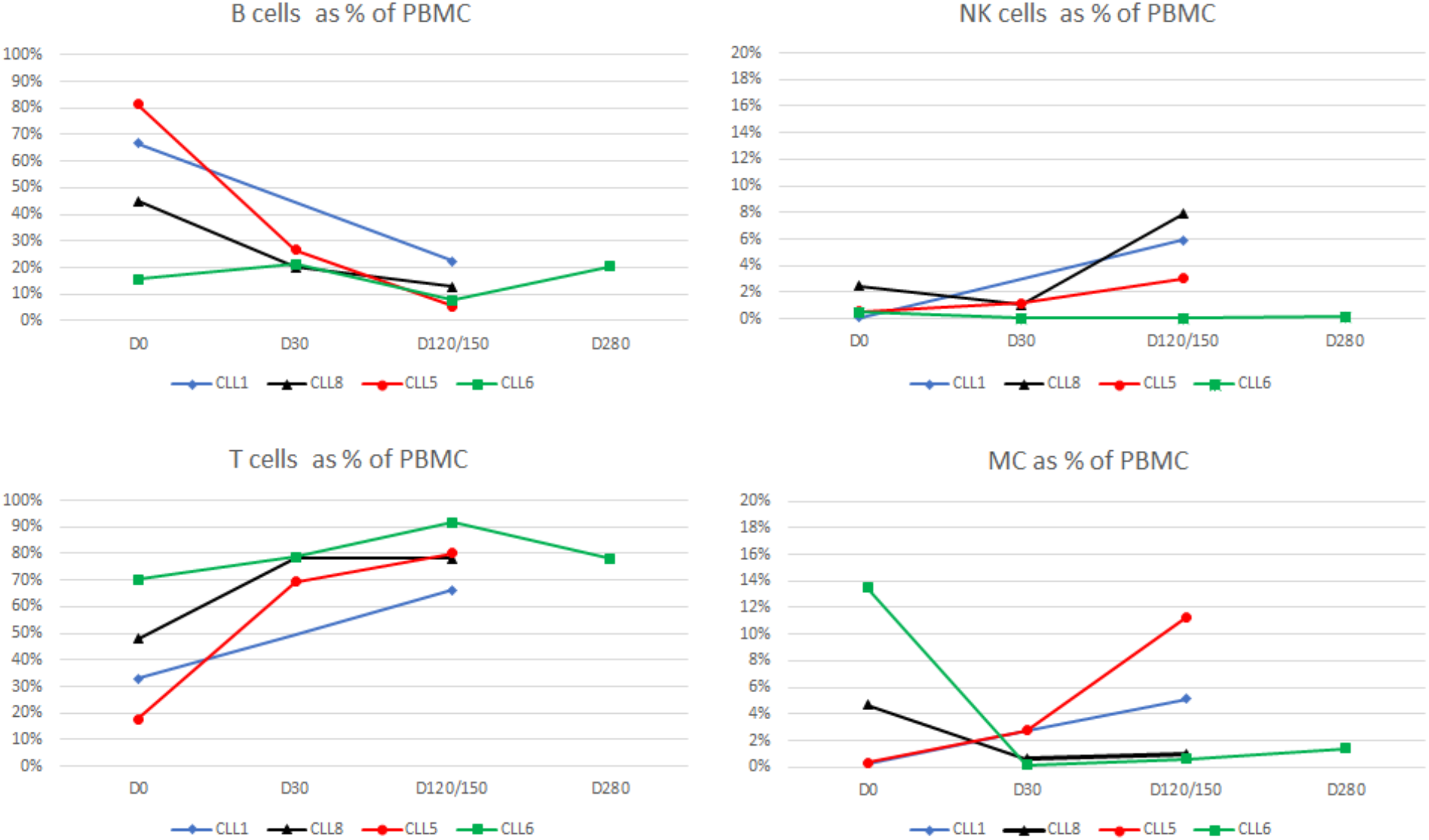
Predicted percentages of BC, TC, NK and MC along the treatment days. Over time, CLL1, CLL5, and CLL8 show the decline of the percentage of cells whose gene expression profiles resemble healthy B cells. CLL1, CLL5, and CLL8 show the increased percentages of cells that look like healthy T cells and NK cells. CLL1 and CLL5 show the increase of the number of cells that look like healthy monocytes while CLL8 shows a decline to the near-zero value. CLL6 shows distinctly different profile changes as compared to the other three patients.

#### *Therapy Day 30* (Figure 6)

Predicted percentages of B cells for CLL5, CLL6, and CLL8 were between 20% and 30%, above normal but lower than on Day 0. Predicted percentages of T cells increased but were below normal range in CLL1 (33%), low-normal in CLL8 (48%), and high-normal in CLL6 (70%). Predicted percentages of monocytes were low in CLL5 (1%), CLL6 (0%) and CLL8 (1%). Predicted percentages of NK cells were low in all patients – 1% in CLL5 and CLL8 and 0% in CLL6. In comparison with Day 0, in all patients the percentages of predicted B cells declined, percentages of T cells increased, while monocytes and NK cells remained low. Day 30 data were not available for CLL1.

#### *Therapy Day 120/150* (Figure 6)

Predicted percentages of B cells further declined in all patients – CLL1 (22%), CLL5 (5%), CLL6 (8%), and CLL8 (13%), while those of T cells increased to normal ranges in CLL1 (66%), CLL8 (78%), and in CLL5 (80%) reached normal or high-normal range, while in CLL6 (92%) exceeded normal range. Predicted percentages of monocytes were below normal healthy ranges in all patients – CLL1 (5%), CLL5 (3%) and CLL6 (0%), and CLL8 (8%). Predicted percentages of NK cells varied – 1% in CLL6 and CLL8, 5% in CLL1 and 11% CLL8. In comparison with Day 0, in all patients the percentages of predicted B cells declined, percentages of T cells increased, while changes in monocytes and NK cells varied.

#### *Other* (Figure 6)

Day 280 in CLL6 had high percentage of B cells (20%), high-normal of T cells (78%), and low of monocytes (1%) and NK cells (0%). No data at this time point are available for other patients.

We compared data of Day 0 and Day 120/150 by radar plots of ANN predictions (Figure 7). The healthy control showed deltoid shape (dashed line in Figure 7). On Day 0, CLL1, CLL8, and CLL5 showed triangular shapes: shift from T cells to B cells and close-to-zero percentage values of monocytes and NK cells. CLL6 showed a completely different shape: similar to healthy PBMC but with zero NK cells. At Day 120/150, ANN predictions of CLL1 and CLL8 were similar to the healthy sample, but with very low monocyte percentage. On Day 120, CLL6 showed very high predictions of T cells (92%), a low number of B cells (8%) and negligible numbers of predicted monocytes and NK cells. On Day 150, CLL5 resembled the healthy profile but with a larger number of T cells.

**Figure 7.**
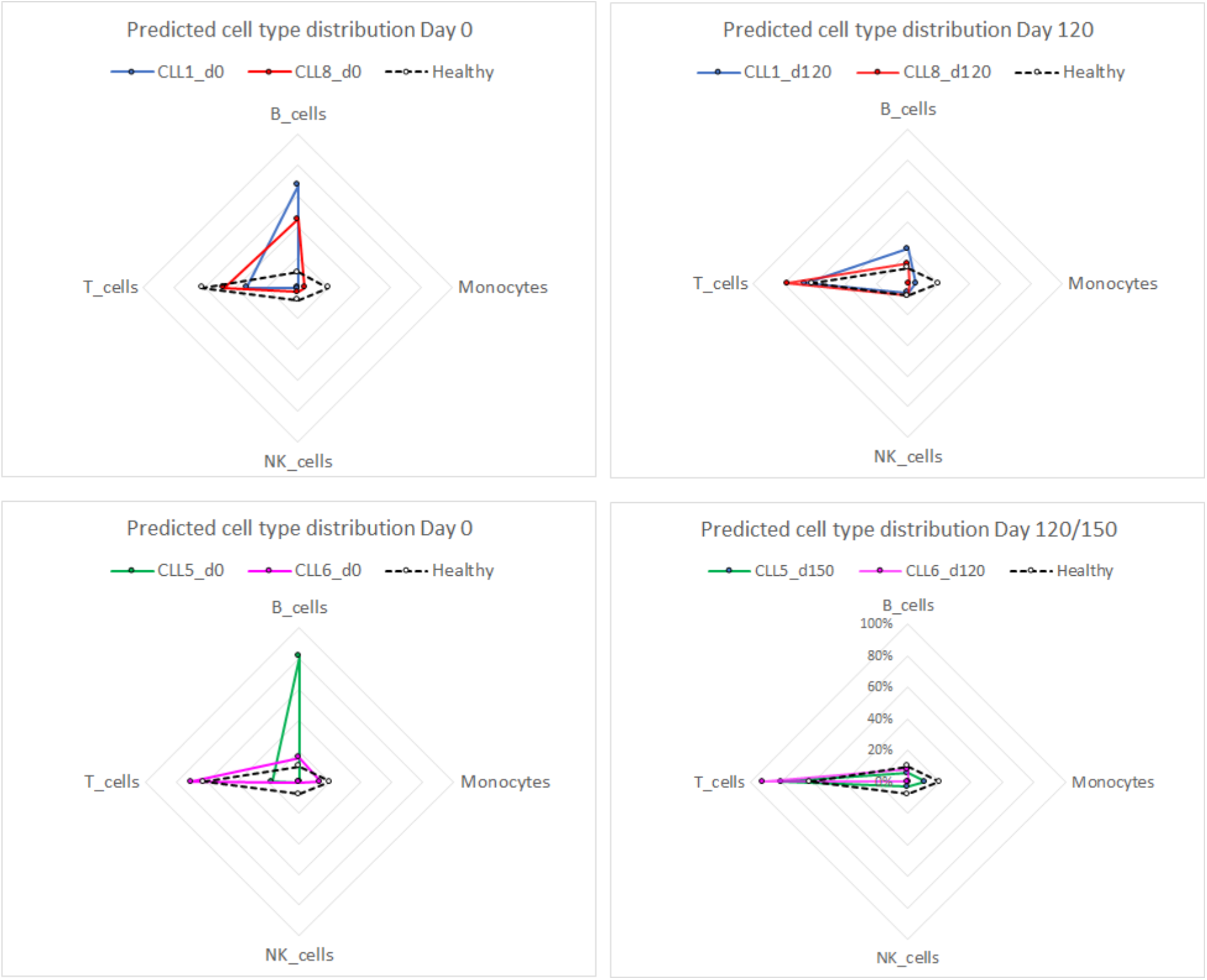
Radar graphs showing predicted percentages of cell types using ANN predictor for healthy PBMC (BC, MC, NK, and TC) from the CLL samples. The graphs show predictions on Day 0 for full remission outcomes (CLL1 and CLL8, upper left); Day 120 CLL1 and CLL8, upper right), Day 0 of relapse (CLL5) and partial remission (CLL6) (lower left); Day 120 of CLL6 and Day 150 of CLL5 (lower right). All patients show profound changes between Day 0 and Day 120/150. CLL1, CLL8 and CLL5 show similar prediction profiles, while CLL6 shows different prediction profile to the other three patients.

### Interpretation of results

We analyzed SCT data sets representing treatment time-points of four CLL patients and one healthy control data set using five comparison tools. These tools provide a comparative analysis between different types of patients: two patients with complete remissions, one patient with partial remission, and one patient with relapse of CLL. The first tool, violin plots, provided the analysis of total counts and their statistical distributions. All pre-treatment distributions had elongated needle-like shapes with high medians and broad IQR ranges. Samples representing complete remission showed a rapid reduction of total counts, as early as on Day 30, resulting in a very low median and very narrow IQR at Day 120. Partial remission samples show little changes from pre-treatment up to Day 120, but low median and IQR at Day 280. The second tool, analysis of expression relative to our standard QC threshold of 670 total counts per cell provided further insight into quantitative changes of gene expression. By Day 120, samples representing total remission had only 10-15% of cells with gene expression counts above the QC threshold. Partial remission sample had this pattern delayed, while the relapse sample at day 120 had 93% of samples with counts above the QC threshold.

The results of the first two tools are consistent with the reported changes in gene expression upon Ibrutinib therapy. These changes involve inhibitory effects across multiple signalling pathways, large reduction of pathway signature scores, and the absence of upregulated biological pathways. It was observed that 76.3% of differentially expressed genes decreased and 23.7% increased their level of expression upon Ibrutinib therapy [37]. It was proposed that Ibrutinib therapy shifts CLL cells into resting state with much of the signalling pathways turned down. Our results indicate that successful shutdown of signalling pathways may be apparent within the first 30 days of the therapy (Figure 4). The rapid decline of median value of total gene expression counts compared to the pre-treatment baseline is a possible early predictor of successful Ibrutinib therapy.

The third tool we used was the hierarchical clustering of all cells in the samples. The healthy control data set shows a large diversity and the presence of numerous micro-clusters. In contrast, the baseline CLL shows a high similarity of groupings of cells by clustering (Figure 5). On Day 120/150, the complete remission samples showed the emergence of healthy-like cluster maps (lower right corner), more pronounced in CLL1, but also emerging in CLL8 and CLL5, while it was not apparent in CLL6. Our hierarchical clustering results suggest that at Day 120 from the start of the therapy, the emergence of the diversity of gene expression resembling those in healthy control data sets is noticeable.

The fourth and the fifth tools focus on the analysis of CLL samples by using healthy PBMC cell type classifier. The classification of cells from CLL samples by ANN does not necessarily predict the cell type in CLL samples but it predicts their similarity to healthy PBMC cell types considering the total gene expression. Overall, CLL1, CLL8, and CLL5 (complete remission and relapse) showed similar trends for B cells, T cells, and NK, but not for the monocytes (Figure 6). CLL6 showed different patterns of PBMC percentage predictions to the other three CLL samples. Interestingly, CLL5 showed the highest increase of monocytes (0.3% to 11.3%) between Day 0 and Day 150. It was found that the high level of circulating monocytes (monocyte-derived cells in CLL environment) is associated with adverse clinical outcomes for CLL patients [38]. This is consistent with our observation that the complete remission patients (CLL1 and CLL8) showed low count (<5%) of cells whose gene expression resembles profiles of healthy monocytes (predicted as monocytes by our ANN) (Figure 6 and 7). The low number of predicted monocytes was maintained through Day 120 of the therapy. The partial remission patient CLL6 showed higher number of monocyte predictions (within the healthy range) on Day 0, and Day 150 the number of predicted monocytes reverted to zero. The patient CLL5 (relapse) had zero monocyte count but the predicted monocytes were similar to the healthy sample on Day 120. The number of predicted NK cells was low in all cases (<3%) on Day 0 and increased in all patients except for the patient with partial remission (CLL6).

## Conclusion and discussion

We developed a set of tools that support supervised machine learning for classification of CLL cells by their similarity to healthy PBMC and the analysis of gene expression changes during Ibrutinib therapy. The distribution of gene expression medians and IQR in individual data sets were calculated. The absolute value of gene expression was compared across different therapy days in individual patients. The similarity of gene expression between individual cells within each sample was studied using ANN classification. Individual cells from PBMC were labelled by ANN by their similarity of gene expression to the profiles of healthy PBMC. The PBMC samples from CLL patients were taken at several treatment timepoints (Days 0, 30, 120/150, and 280), as available. After each cell in individual CLL samples was labelled, the proportions of BC, MC, NK, and TC were calculated. We observed specific patterns that show consistency with the results of previous clinical studies.

We observed several patterns in gene expression changes in CLL patients change during the ibrutinib therapy:

- Gene expression patterns in CLL patients before Ibrutinib therapy show high median and broad interquartile ranges. Patients with complete remission within three years showed rapid drop of both median gene expression and the corresponding IQR. The patient with relapse showed lower median gene expression and IQR than other patients at Day 0. The same patient showed an increase in median gene expression and IQR at Day 30, and reduction of both terms after Day 150. The patient with partial remission showed marked reduction of median gene expression and IQR by Day 280 of therapy. A clear pattern of rapid decline of both the median gene expression and IQR was seen in patients with the full remission of CLL (Figure 3).
- The intensity of gene expression, measured as the percentage of cells in samples that have more or equal to 670 total counts is low in samples CLL1 and CLL8, associated with positive outcomes in this study (Figure 4). Persistent high intensity of gene expression was observed on Day 0 and day 120/150. These observations are consistent with the results published in [37]. Ibrutinib therapy shows strong inhibitory effect on multiple molecular pathways resulting in broad gene expression changes in CLL cells. Ibrutinib induces “silencing” of cancer cells and the cancer microenvironment bringing cancer to resting-state followed by the attrition of residual cancer over time [37]. It was reported that as early as after two days of therapy Ibrutinib shuts down ongoing inflammatory responses in peripheral blood and lymph nodes [40]. The “silencing” pattern was observed in CLL1 and CLL8 samples, was not observed in the relapse sample CLL5, and was delayed in the partial remission sample CLL6 (Figure 4).
- Hierarchical clustering of all cells has shown a high level of similarity between cells within each sample on therapy Day 0 and low diversity of clusters. On day 120 the sample CLL1 showed clear emergence of healthy-like cell diversity (lower right corner of CLL1 Day 120). Another full remission sample CLL8 showed similar emergence of cluster diversity (lower right corner of CLL8 Day 120), but it was less pronounced than in CLL1, consistent with the observed level of lymphocytosis – moderate in CLL1 and weak in CLL8 (Supplemental Table 1). We hypothesise that the increase in the number of lymphocytes represents the lymphocytes that show gene expression profiles similar to healthy cells.
- The analysis of similarity of gene expression profiles between Ibrutinib treatment samples and healthy control PBMC by ANN (similarity of CLL PBMC to healthy cells) showed similar profiles in patients with complete remission (CLL1 and CLL8): a large proportion of predicted B-cells, reduced proportions of predicted T cells and minimal numbers of predicted NK cells and monocytes at Day 0. At Day 120, the predicted profiles shifted to healthy ranges (as a percent of the total PBMC) of T cells, slightly elevated (relative to healthy ranges) numbers of B cells, close to zero predicted monocytes, and low-normal range of predicted NK cells (Figure 7). The relapse case CLL5 showed similarity to CLL1 and CLL8 at Day 0, but similarity to healthy PBMC profile at Day 120. The partial remission sample CLL6 showed predicted T cells in healthy range, predicted B cells slightly elevated healthy range, predicted monocytes within the healthy range, and no predicted NK cells. At day 150, predicted T cells were 93% with the rest of predicted cells labelled as B cells. Taken together, the predictor of healthy PBMC cell types showed that patterns in CLL1 and CLL8 samples (successful therapy) were similar, while partial remission sample CLL6 and relapsed sample CLL5 showed different patterns. We note that the analysis of common cell type markers indicates that actual cell type in CLL and predicted cell types by ANN healthy PBMC classifier are not necessarily the same. PBMC in CLL are highly dysregulated and many of the cells display markers of multiple cell types (data not shown). In this analysis we only look at overall similarity of predicted cell types in comparison with healthy PBMC (comparison of proportions) but did not attempt to assign the actual cell type.

Overall, the results of this study suggest that patients that experience full remission show early and large reduction in median gene expression level, their IQR, and the majority of actual counts reduced by 5-fold or more mostly below the usual QC level for single cell analysis (Figure 3 and 4). This is combined with the increase of the diversity of cell clusters (comparable to healthy cluster maps) and reduction of CLL-like clusters and gradual restoration of cells that resemble healthy PBMC (Fig. 5, 6 and 7). While the number of CLL patients in this study is insufficient for strong conclusion, we suggest the possibility of early prediction of responders by observing the patterns of gene expression consistent with the rapid shutdown of molecular pathways by Ibrutinib. We propose new single cell analytical tools that use combination of clustering and supervised classification to predict early responders in Ibrutinib therapy. A prognostic scoring system for survival of CLL patients treated with ibrutinib was reported [40]. This prognostic system uses four independent variables (aberration of TP53, relapsed CLL, B2 microglobulin concentration ≥ 5 mg/L, and lactate dehydrogenase > 250 Units/L) to stratify patients into three risk groups: high, intermediate, and low risk. We propose a model of early response to Ibrutinib for an early response associated with complete remission, based on the analysis of single cell gene expression counts (Figure 8) and supervised Machine learning (Figures 6 and 7). This model needs to be validated with additional data and further studies. Furthermore, gene expression in CLL before and during therapy appears to reflect patterns of dysregulation in PBMC samples. This raises an attractive hypothesis that the type of responses to Ibrutinib may be predicted before the therapy. This is subject of our ongoing research.

**Figure 8.**
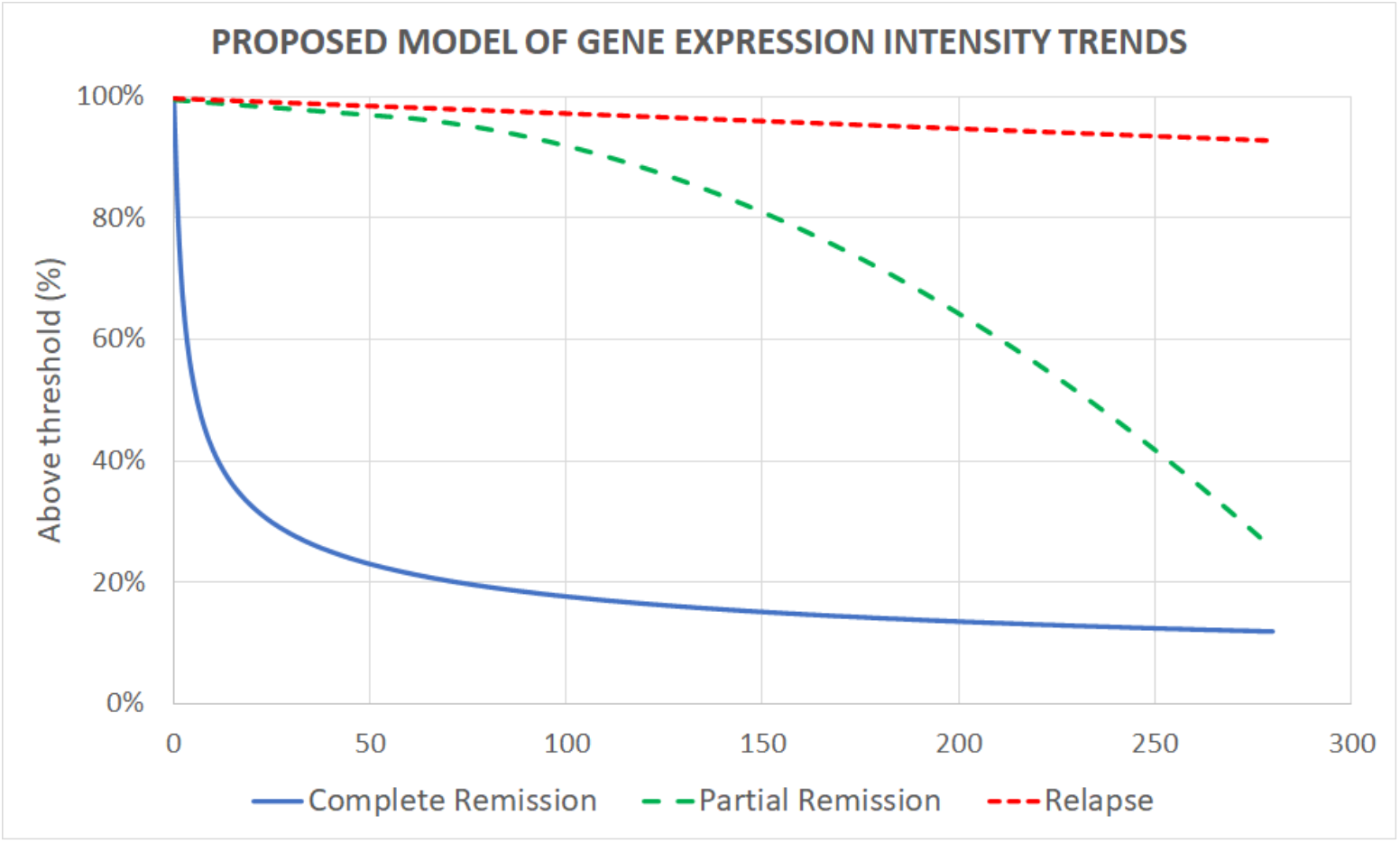
The model of response to Ibrutinib based on clinical observations [33,35,37-40] and the results of the analysis in this study. Gene expression counts from single cell samples in patients with complete remission within 3 years show rapid drop of total gene counts within the first few days of the therapy. The samples from partial remission patients show little changes at the beginning, followed by gradual reduction of total gene counts in CLL PBMC. Relapsed CLL cases do not show notable decline of the total gene counts. This proposed model needs further validation and refinement.

## Abbreviations

SCT: Single Cell Transcriptomics
ANN: Artificial neural network
BC: B cells
CLL: chronic lymphocytic leukemia
CLL1: chronic lymphocytic leukemia patient 1
CLL5: chronic lymphocytic leukemia patient 5
CLL6: chronic lymphocytic leukemia patient 6
CLL8: chronic lymphocytic leukemia patient 8
DC: dendritic cells
NK: natural killer cells
PBMC: peripheral blood mononuclear cells
PR: partial remission (of CLL)
QC: quality control
RE: relapse (of CLL)
TC: T cells

## Acknowledgements

The cost of publication of this article was supported by the Sensors, Sensor Networks and Instrumentation Research Group, at the Faculty of Science and Engineering, University of Nottingham Ningbo China.

## Authors’ contribution

VB, DK, and ZC conceived the idea. VB designed the study. ML, XL and VB performed data preparation, tool development, programming, and analysis. VB, ML, AB, DK, HJ, and ZC discussed the results. All authors contributed to writing the paper. ML and VB took the lead in writing the manuscript.

## Availability of data and materials

All datasets from this study are available at http://projects.met-hilab.org/SCTdata/CLL001

## Ethics approval and consent to participate

Ethics approval was not required for this study.

## Supplementary information

Supplemental Table 1. Metadata describing samples as described by the sources.

Supplemental Table 2. The results of basic statistical analysis of the data sets.

